# Histone lactylation antagonizes senescence and skeletal muscle aging via facilitating gene expression reprogramming

**DOI:** 10.1101/2023.05.26.542348

**Authors:** Xuebin Zhang, Fanju Meng, Wencong Lyu, Jianuo He, Ran Wei, Zhehao Du, Chao Zhang, Yiting Guan, Xiaoke Huang, Guoliang Lyu, Xiao-Li Tian, Lijun Zhang, Wei Tao

**Affiliations:** The MOE Key Laboratory of Cell Proliferation and Differentiation, School of Life Sciences, Peking University, Beijing 100871, China; Department of Human Population Genetics, Human Aging Research Institute (HARI) and School of Life Sciences, Nanchang University, Nanchang 330031, China

**Author notes:** Correspondence Wei Tao,. Lijun Zhang,. These authors contributed equally to this work (Xuebin Zhang, Fanju Meng, Wencong Lyu). Funding information The National Key Research and Development Project, Grant No. 2021YFA0909300; The National Natural Science Foundation of China, Grant No. 32241006.

**Keywords:** Histone lactylation, Senescence, Skeletal muscle aging, Metabolism

## Abstract

One of the prominent drivers of cellular senescence and/or aging is epigenetic alteration, through which orchestrated regulation of gene expression is achieved during the processes. Accumulating endeavors have been devoted to identifying histone modifications-related mechanisms underlying senescence and aging. Here, we show that histone lactylation, a recently identified histone modification bridging metabolism, epigenetic regulation of gene expression and cellular activities in response to internal and external cues, plays a crucial role in counteracting senescence as well as mitigating dysfunctions of skeletal muscle in aged mice. Mechanistically, the abundance of histone lactylation is markedly decreased during senescence and aging but restored following manipulation of the metabolic environment. Genome-wide distribution profiling and gene expression network analysis uncover that the maintenance of histone lactylation level is critical for suppressing senescence and aging programs via targeting of proliferation- and homeostasis-related pathways. We also confirmed that the level of histone lactylation is not only controlled by glycolysis but also regulated by NAD^+^ content *in vivo*. More intriguingly, running exercise enhances the level of histone lactylation and reconstructs the cell landscape and communications of mouse skeletal muscle, leading to rejuvenation and functional improvement. Our study highlights the role of histone lactylation in regulating senescence as well as aging-related tissue function, implying that this modification could be used as a novel marker of senescence, and provides a potential target for aging intervention via metabolic manipulation.

## Main

Senescence, which is characterized by permanent cell cycle arrest along with increased expression of cyclin-dependent kinase inhibitors and the senescence-associated secretory phenotypes (SASP), can be triggered by a variety of intrinsic and extrinsic environmental cues and developmental signals^1^. It is widely accepted that senescence plays important roles in a variety of physiological and pathological processes^2^. One of the hallmarks as well as drivers of senescence and/or aging, epigenetic alteration, in particular the rearrangement of histone modifications, is key to the progression of senescence via modulating the gene expression network, chromatin architecture organization and genome stability maintenance^3–7^. A plethora of studies have substantiated the promising applications of small molecules that could affect epigenetic information in life span expansion and aging intervention^8–11^.

Notably, senescent cells or aged tissues still maintain highly metabolic activities and cellular metabolic status is crucial to not only cell proliferation but also other cell fate-determining processes, during which epigenetic regulation of gene expression is also required^12^. Metabolic intermediates possess the chemical groups required for the post-translational modification (PTM) of histones and non-histone proteins^13^; for example, acetyl-CoA produced in the mitochondria provides acetyl groups for histone acetylation. The discovery of such modifications has greatly expanded our understanding of the multiple roles of histone codes in the regulation of chromatin state, gene expression, and cell phenotype^14^.

Histone lysine lactylation, especially histone 3 lysine 18 (H3K18la) is a novel histone modification with functions that have not yet been extensively explored. To date, it has been proven that histone lactylation plays a pivotal role in gene expression regulation and is critical to the metabolism–epigenetics axis in modulating cellular activities, e.g., the immune response of macrophages and oncogenesis^15^. Histone lactylation is also reported, though not in much detail, to be associated with other biological processes, including tissue repair, proliferation, metabolism, neurodevelopment, and embryonic development^15–24^. These associations not only extend its potential physiological functions but also further imply that lactate mainly generated from glycolysis is not merely a metabolic waste but instead a key regulator of cell fate because it provides lactyl groups for lactylation of histones. The level of histone lactylation is closely related to metabolic kinetics in response to cellular activities and glycolytic enzymes, acyltransferase (P300), and deacylase (HDAC1/3), which may contribute to writing, reading or erasing histone lactylation with recruitment of cofactors^15, 25, 26^. Although histone lactylation has been shown to occur in both promoters and enhancers to activate the expression of target genes^15, 19, 27^, its role and mechanisms in regulating senescence and tissue functions during aging remain elusive.

In this study, using a variety of techniques, models, and histone sites, we explored the characteristics and functions of histone lactylation in senescence and aging and showed that the abundance of histone lactylation declines markedly during replicative senescence in human embryonic lung fibroblasts (IMR90 cells), mouse embryonic fibroblasts (MEFs), and human umbilical vein endothelial cells (HUVECs). Taking IMR90 cells as the main *in vitro* model, we demonstrated that both hypoxia exposure and NAD^+^ resumption delay senescence, which is accompanied by an increase in the level of histone lactylation. Enrichment of histone lactylation in promoters of anti-aging genes related to cell proliferation and homeostasis antagonizes senescence, while histone lactylation-targeted enhancers can regulate cell morphogenesis. Furthermore, we demonstrate that the abundance of histone lactylation in mouse skeletal muscle is increased after running exercise, leading to remarkable reconstruction of cell composition, and subsequently rewiring cell communication networks. The cell rearrangement events in skeletal muscle indicate that high histone lactylation levels may reflect a younger physiological state and better organ function. Overall, our data reveal the role of histone lactylation and could suggest it serving as a candidate marker to assess senescence and the aging state.

## Results

### Histone lactylation level declines during senescence

The discovery of histone lactylation has provided novel insights into the role of the metabolism–epigenetic axis in the process of senescence. To reveal the relationship between histone lactylation and senescence, we constructed three replicative senescence models: human embryonic lung fibroblasts (IMR90 cells), mouse fibroblasts (MEFs, and human umbilical vein endothelial cells (HUVECs). At the beginning of passage, cells grow rapidly, which is defined as the young state; by contrast, cells cease to proliferate when confluence is reached, which is referred to as the senescent state.

Multiple markers were used to confirm the senescence state, i.e., positive SA-β-gal staining, loss of Ki-67 expression, decreased protein expression of Lamin B1, and increased protein expression of p16 and p21. Consistently, the percentage of cells with SA-β-gal^+^ staining was significantly increased (Additional file: Fig. S1A–C) and Ki-67 expression was abolished (Additional file: Fig. S1D–F) in the three models of senescence. Additionally, Lamin B1 expression was decreased, and p16/p21 expression was increased. Moreover, the protein levels of histone lactylation at different H3 sites, including H3K9la, H3K14la, and H3K18la, were markedly decreased (Fig. 1A–C), suggesting that histone lactylation declines globally during senescence across cell types.

**Fig. 1.**
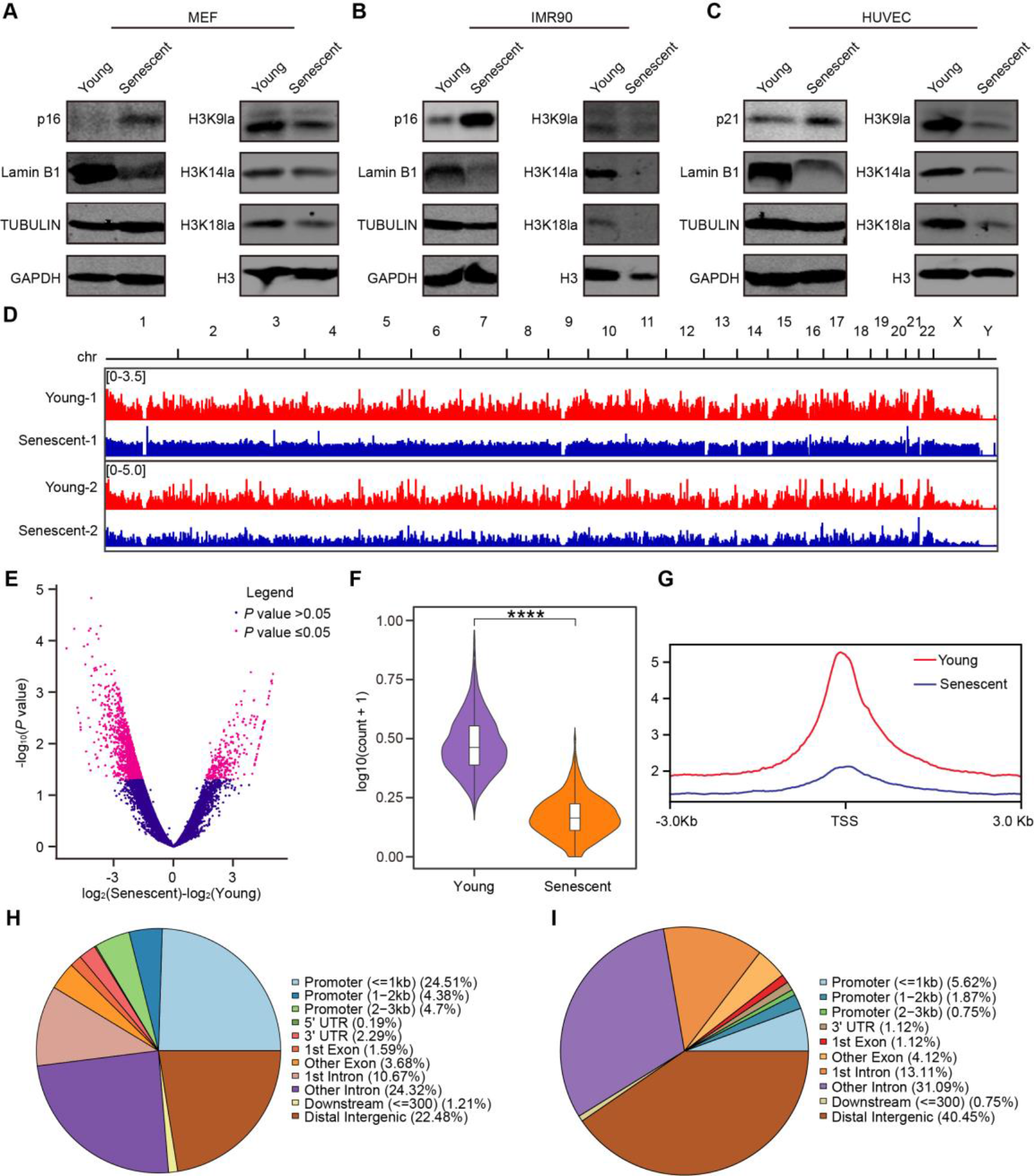
**The level of histone lactylation decreases in senescent cells. A–C**, Immunoblotting of Lamin B1, p16 (**A, B**), p21 (**C**), H3K9la, H3K14la, H3K18la, and H4K12la in young and senescent MEF cells (**A**), IMR90 cells (**B**), and HUVECs (**C**). TUBULIN, GAPDH, and H3 served as the loading controls. **D**, Snapshot showing the H3K9la abundance at the whole-genome level during aging. Two biological repeats were used to perform CUT&Tag-seq. **E**. Volcano plot of the differential H3K9la peaks in IMR90 cells during senescence. The x-axis represents the fold change in peak signals between young and senescent samples, and the y-axis represents the *P* value with the significance threshold set to 0.05. **F**, Violin plot of the signal intensity of the downregulated H3K9la peaks in IMR90 cells during senescence. \*\*\*\**P* < 0.0001. **G**, Profiles of H3K9la peak signals in the range of 3 kb upstream and 3 kb downstream of TSSs throughout the genome in young and senescent IMR90 cells. **H**, Pie chart of the proportional distribution of downregulated H3K9la peaks in genomic elements in IMR90 cells during senescence. **I**, Pie chart of the proportional distribution of upregulated H3K9la peaks in genomic elements in IMR90 cells during senescence.

Next, to further confirm whether histone lactylation is decreased at the genome-wide level, we performed the CUT&Tag-seq for H3K9la in IMR90 cell. The abundance of histone lactylation was significantly decreased throughout the entire genome, which is consistent with the immunoblotting data (Fig. 1D). Then, we counted the differential peaks of H3K9la, of which the total numbers in senescent cells were remarkably lower than those in young cells (Fig. 1E). When we focused on the decreased differential peaks of H3K9la during senescence, the significant downward trend caught our attention (Fig. 1F). In particular, the drop in histone lactylation was even more pronounced in the promoter region than in other regions (Fig. 1G and Additional file: Fig. S2). Statistical analysis of the sequencing data showed that the decreased H3K9la peaks during senescence were mainly distributed in promoters, distal intergenic regions, and introns (Fig. 1H). Compared with the downregulated H3K9la peaks in promoters, we found that the upregulated H3K9la peaks were mainly distributed in distal intergenic regions during senescence (Fig. 1I).

Then, the lactate levels in young and senescent cells were measured in view that the histone lactylation is derived from the metabolite lactate. Unexpectedly, there was no association between the levels of endogenous lactate and histone lactylation during senescence (Additional file: Fig. S3), which is consistent with previous reports^27^. These results demonstrated the uncoupling between lactate level and histone lactylation during senescence. Nonetheless, when exogenous sodium lactate was applied to MEFs and HUVECs, the histone lactylation level increased, and the senescence of MEF and HUVEC was delayed (Additional file: Fig. S4A–D), indicating that additional lactate remarkably increased histone lactylation levels, in accordance with the results of previous studies^15, 20^. The results of bulk RNA-seq and subsequent Gene Ontology (GO) enrichment analysis showed that anti-aging pathways including proteolysis, Wnt signaling, and extracellular matrix organization, were upregulated due to sodium lactate (NALA) treatment (Additional file: Fig. S4E, F), and pro-aging pathways, such as reactive oxygen species (ROS) production, were decreased (Additional file: Fig. S4G). These results also reveal that the transcription program of senescence can be reversed by exogenous lactate-increased histone lactylation.

### The modulation of metabolism is capable of manipulating histone lactylation abundance

Considering that exogenous lactate could induce restoration of histone lactylation and further inhibit the senescence process, we hypothesized that active intervention of lactate level in cells is sufficient to regulate the abundance of histone lactylation and influence the senescence fate. In addition, due to the long process and the complexity of regulatory factors for senescence, we deemed it worthwhile to find an instantaneously sensitive and stable system with which to explore the regulatory axis of histone lactylation in senescence.

Since hypoxia can produce large amounts of endogenous lactate^28^, and previous researches evince that hypoxia considerably extends cell life span^29^, we established a physiological hypoxia (3% O2) model in IMR90 and MEF cells to verify the hypothesis that whether lactate alteration via external environment manipulation is able to regulate the level of histone lactylation and in turn intervene senescence. The proliferative capacity of IMR90 cells under normoxia was very poor and they reached senescence rapidly; however, following transfer to hypoxia, the proliferation potential of IMR90 cells was activated, and the process of senescence was significantly delayed (Fig. 2A, B). The same phenomenon was also observed in MEFs (Additional file: Fig. S5A).

**Fig. 2.**
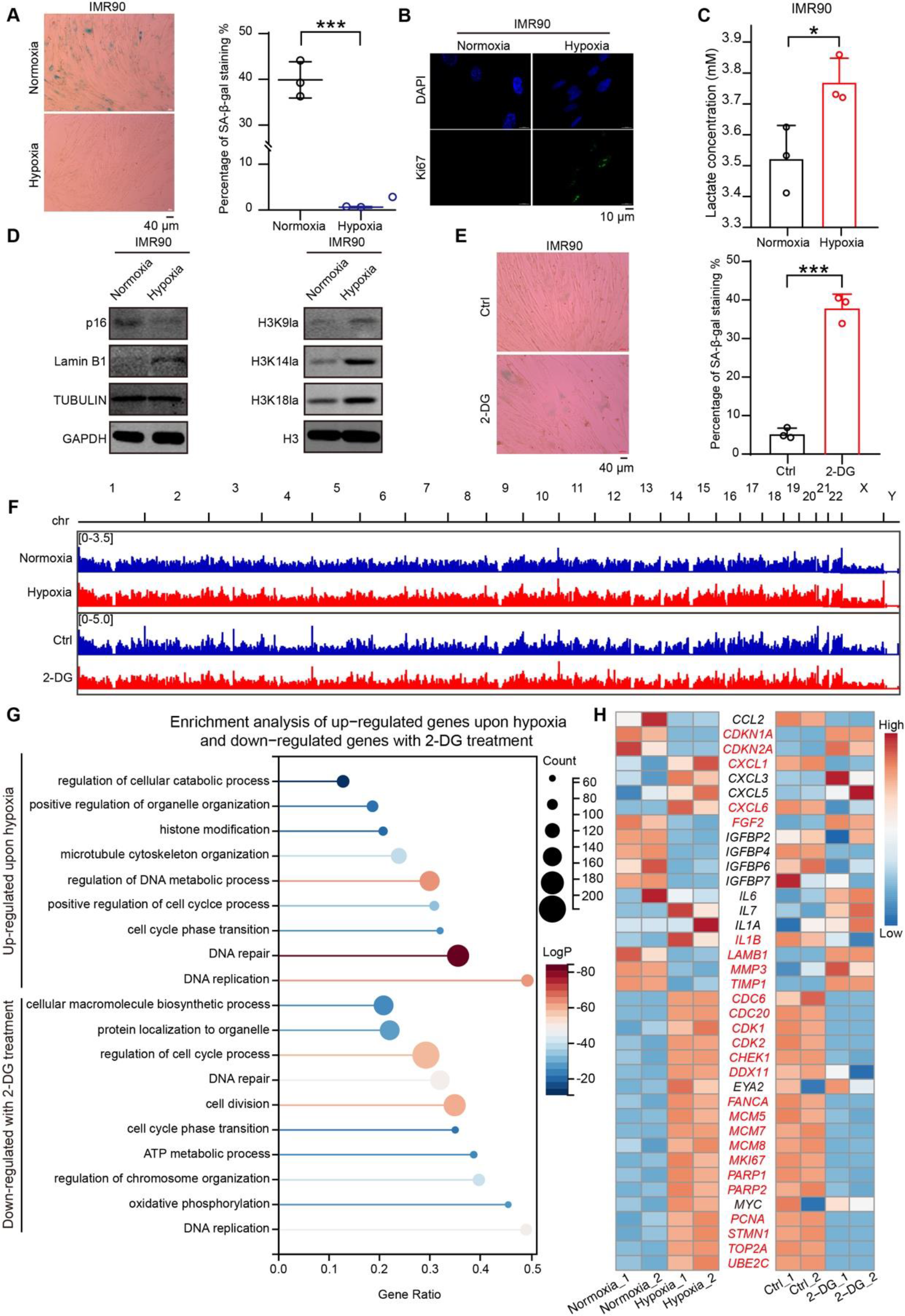
**Hypoxic environment restores the loss of histone lactylation during senescence. A**, SA-β-gal staining of IMR90 cells cultured under normoxia or hypoxia. The percentages of SA- β-gal^+^ cells are shown on the right. **B**, Immunofluorescence of Ki67 and DAPI in IMR90 cells cultured under normoxia or hypoxia. **C**, Intracellular lactate levels in IMR90 cells cultured under normoxia or hypoxia. **D**, Protein levels of p16, Lamin B1, H3K9la, H3K14la, and H3K18la in IMR90 cells cultured under normoxia or hypoxia. TUBULIN, GAPDH, and H3 served as the loading controls. **E**, SA-β-gal staining of IMR90 cells with or without 2-DG treatment in a hypoxic environment. The percentages of SA-β-gal^+^ cells are shown on the right. **F**. Snapshot showing the H3K9la abundance at the whole-genome level with hypoxia or 2-DG treatment. **G**, Functional enrichment analysis of genes that were upregulated under hypoxia or downregulated with 2-DG treatment in IMR90 cells. **H**, Heatmap showing the example genes upregulated under hypoxia or downregulated with 2-DG treatment in IMR90 cells. Genes written in red letters represent the overlapping genes in the two treatments.

Additionally, the lactate level was significantly increased in IMR90 cells under hypoxia compared with that under normoxia (Fig. 2C). Under hypoxia, the protein expression level of the cell cycle suppressor p16 was downregulated, while that of Lamin B1 was up-regulated (Fig. 2D). In addition to a slowing of the senescence process, marked increases in histone lactylation abundance were observed, including increases in H3K9la, H3K14la, and H3K18la (Fig. 2D). Consistently, the protein expression level of Lamin B1 and the abundance of histone lactylation were increased in MEFs (Additional file: Fig. S5B). These results suggested that hypoxia recovers histone lactylation via enhancing intracellular glycolysis flux.

However, the retarding effect of hypoxia on senescence was mitigated when IMR90 cells cultured in a hypoxic environment were treated with a glycolysis inhibitor 2-DG (2-Deoxy-D-glucose) (Fig. 2E). In addition, the abundance of histone lactylation was decreased at the genome-wide level (Fig. 2F). The content of histone lactylation was periodically controlled by hypoxia and 2-DG exposure (Additional file: Fig. S6). To further confirm that the fluctuation of glycolysis flux directly influences the histone lactylation level, and the histone lactylation abundance also indicates the cell proliferative state, bulk RNA-seq analysis upon hypoxia exposure and 2-DG treatment was performed. We found that cells exposed to a hypoxic environment displayed up- regulated proliferation-related pathways, e.g., DNA repair, DNA replication, and DNA metabolism. By contrast, these pathways were down-regulated correspondingly upon 2-DG addition (Fig. 2G). The gene expression profile also changed periodically, and a large number of genes regulating senescence were differentially expressed under hypoxia and 2-DG treatment, e.g., *CDKN1A*, *CDKN2A*, *CXCL1/6*, *FGF2*, *IL1B*, *LAMB1*, *MMP3*, *TMP1*, *CDC6/20*, *CDK1/2*, *CHEK1*, *DDX11*, *FANCA*, *MCM5/7/8*, *MKI67*, *PARP1/2*, *PCNA*, *STMN1*, *TOP2A*, and *UBE2C* (Fig. 2H). These results indicated that histone lactylation mediated the anti-aging effect of physiological hypoxia and the modulation of metabolic environment is an effective means to intervene senescence.

### Histone lactylation reactivates the homeostasis and proliferation to delay senescence progression

The antithetic results between hypoxia and 2-DG treatment suggested that the regulation of histone lactylation on cell proliferation is key to antagonizing senescence. To further demonstrate this, we treated cancer cells (A549, PC3, and U2OS cells) with 2-DG. In accordance with the reported proliferation-related function of histone lactylation^20^, the levels of histone lactylation in A549, PC3, and U2OS cancer cells were decreased following cell cycle blockade with the glycolysis inhibitor 2-DG (Additional file: Fig. S7). The close correlation between histone lactylation and proliferation encouraged us to explore the roles of this modification in senescence.

Then, in-depth analysis of the CUT&Tag-seq data of H3K9la was performed to search for other functions affecting senescence that might be regulated by histone lactylation. We found that the increased H3K9la peaks under hypoxia were mainly distributed in promoters, distal intergenic regions, and introns (Fig. 3A), which was similar to the replicative senescence profile (Fig. 1H). Correspondingly, the distribution characteristics of decreased H3K9la peaks upon 2-DG treatment resembled those increased in the hypoxic environment (Fig. 3B). Especially in the promoter region, histone lactylation showed the opposite trends under the two conditions (Fig. 3C, D). These results indicate that the promoter is the key genomic functional region of histone lactylation in senescence regulation. Meanwhile, previous studies in human M1 macrophages have also demonstrated that histone lactylation activates promoters^15^.

**Fig. 3.**
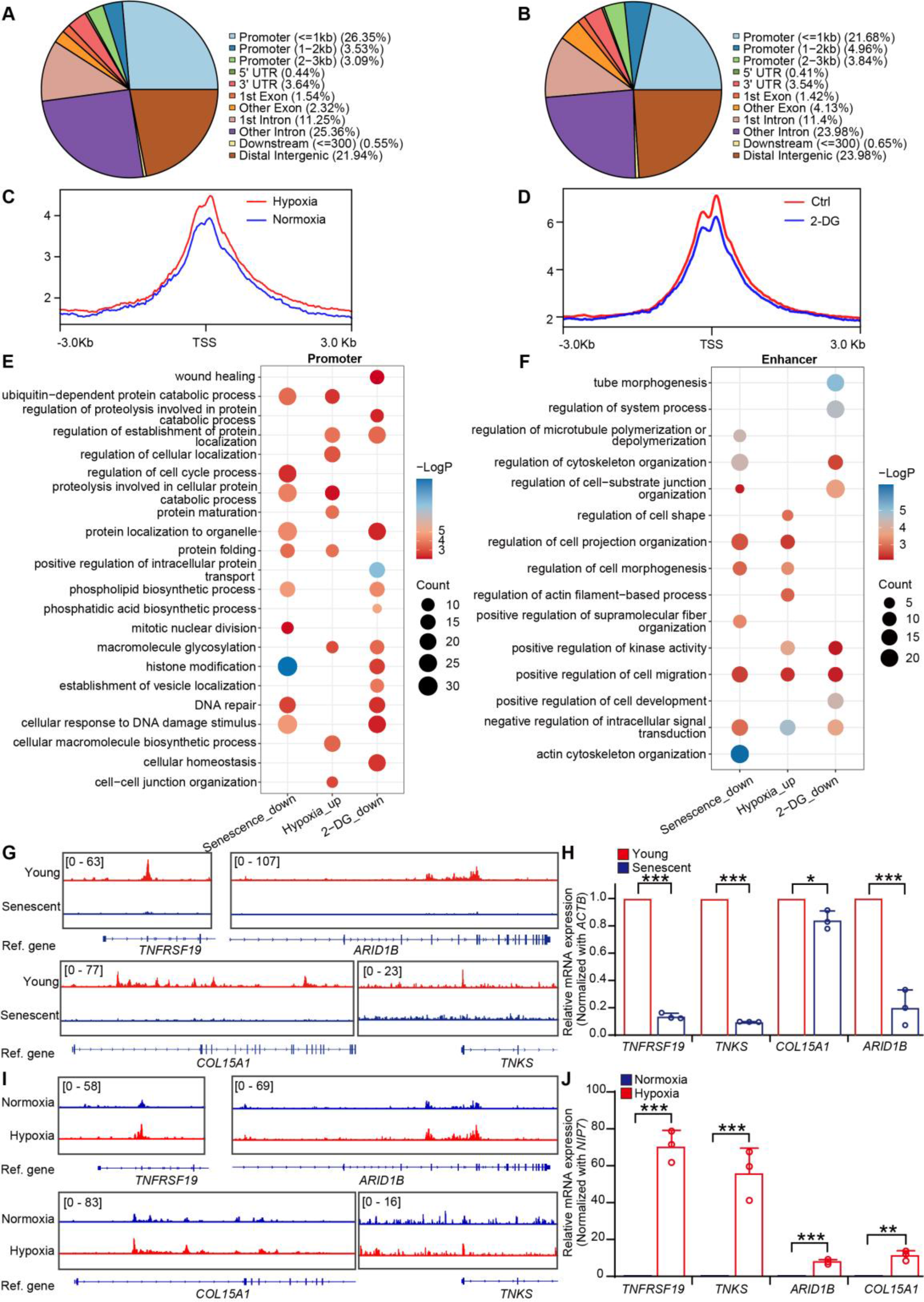
**Increased histone lactylation reprograms the gene expression profile to a youthful state. A**, Distribution of the upregulated H3K9la peaks in genomic elements in IMR90 cells under hypoxia. **B**, Distribution of the downregulated H3K9la peaks in genomic elements in IMR90 cells treated with 2-DG in a hypoxic environment. **C**, Profiles of H3K9la peak signals in the range of 3 kb upstream and 3 kb downstream of TSSs throughout the genome in IMR90 cells cultured under normoxia or hypoxia. **D**, Profiles of H3K9la peak signals in the range of 3 kb upstream and 3 kb downstream of TSSs throughout the genome with or without 2-DG treatment in IMR90 cells cultured in a hypoxic environment. **E**, Functional enrichment analysis of genes near H3K9la-targeted promoters that were downregulated during senescence, upregulated upon hypoxia, and downregulated with 2-DG treatment in IMR90 cells. **F**, Functional enrichment analysis of genes near H3K9la-targeted enhancers that were downregulated during senescence, upregulated upon hypoxia, and downregulated with 2-DG treatment in IMR90 cells. **G**, Snapshots of H3K9la peak signals near *TNFRSF19*, *TNKS*, *COL15A1*, and *ARID1B* in young or senescent IMR90 cells. **H**, mRNA levels of *TNFRSF19*, *TNKS*, *COL15A1*, and *ARID1B* in young or senescent IMR90 cells. The cycle threshold (Ct) values of these genes were normalized to that of *ACTB*. **I**, Snapshots of H3K9la peak signals near *TNFRSF19*, *TNKS*, *COL15A1*, and *ARID1B* in IMR90 cells cultured under normoxia or hypoxia. **J**, mRNA levels of *TNFRSF19*, *TNKS*, *COL15A1*, and *ARID1B* in IMR90 cells cultured under normoxia or hypoxia. The Ct values of these genes were normalized to that of *NIP7*. The error bars represent the S.D. of three independent experiments. Two-tailed, unpaired Student’s *t*-tests were performed. \**P* < 0.05, \*\**P* < 0.01, \*\*\**P* < 0.001. Normoxia, 20% O_2_; Hypoxia, 3% O_2_; TSSs, transcription start sites.

Hence, we extracted the H3K9la differential peaks enriched in the promoter regions for functional enrichment analysis. A more complete picture of histone lactylation-targeted functions came to the surface, comparing the enriched pathways among senescence, hypoxia exposure, and 2-DG treatment. Pathways regulating protein homeostasis, including proteolysis and protein localization, were downregulated in senescence but reactivated in the hypoxic environment. Then, these protein homeostasis pathways were inhibited again with 2-DG treatment (Fig. 3E). The above findings were also substantiated by the down-regulated gene expression profile and the enriched signaling pathways of proteolysis, ribosome biogenesis, DNA metabolism, and cell cycle in IMR90 cells during senescence (Additional file: Fig. S8A). These results suggest that histone lactylation regulates the senescence progression mainly by managing cell proliferation and homeostasis.

Regarding the distribution features of histone lactylation in the genome, the distal intergenic regions also attracted our attention. In view that the distal intergenic regions usually contain enhancers, we analyzed the potential functions of histone lactylation- targeted enhancers. The genes near H3K9la-specific enhancers were involved in response to external signals and cell morphogenesis during senescence, for instance, signal transduction, cell migration, and cytoskeleton organization. Similar to those in the promoter analysis for promoter, the conclusions regarding the functions of histone lactylation-targeted enhancers were supported by forward and reverse experiments (Fig. 3F). These results extend understanding of the roles of histone lactylation in enhancers compared with promoters and suggest that these histone lactylation-targeted enhancers mainly regulate cell morphogenesis.

Subsequently, the differentially expressed genes, *TNFRSF19*, *TNKS*, *COL15A1*, and *ARID1B*, were shown to verify the anti-aging role of histone lactylation in light of that TNFRSF19 may prevent senescence by inhibiting TGF-β signaling^30^; TNKS may promote proliferation by positively regulating Wnt signaling, and loss of TNKS impairs longevity^31, 32^; type XV collagen, which is encoded by *COL15A1*, is important for the function of young tissue^33–35^; and ARID1B is responsible for the maintenance of pluripotency^36^. Indeed, the histone lactylation abundance and the expression levels of these genes were both decreased during senescence (Fig. 3G, H, and Additional file: Fig. S8B). Correspondingly, H3K9la enrichment in the promoters of *TNFRSF19*, *TNKS*, *COL15A1*, and *ARID1B* were restored (Fig. 3I and Additional file: Fig. S8C), and the expression levels of H3K9la-targeted genes were also increased upon hypoxia (Fig. 3J). These data indicated that histone lactylation regulates the expression of anti-aging genes, thereby influencing the progression of senescence.

### The level of histone lactylation depends on NAD^+^ content *in vivo*

Remodeling of the cell metabolic network and the decrease of NAD^+^ content are very distinct features during senescence and aging. NAD^+^ can be reduced to NADH in the oxidation of glyceraldehyde 3-phosphate, and the production of lactate depends on the transition between NAD^+^ and NADH. We hypothesized that the reduction in NAD^+^ content also contributed to the decline in histone lactylation in senescent cells.

Thus, we treated IMR90 cells using NMN (Nicotinamide mononucleotide), a precursor of NAD^+^, which has been shown to slow aging and aging-associated diseases, to further elucidate the anti-aging role of NAD^+^ and histone lactylation^37, 38^. NMN treatment led to restoration of the NAD^+^ level and decreased SA-β-gal signals in senescent cells (Fig. 4A). In the meantime, the expression of senescence markers p16 and p21 was downregulated, and that of Lamin B1 was upregulated (Fig. 4B). Strikingly, when the NAD^+^ content was increased due to NMN addition, the histone lactylation level was also recovered in a NAD^+^ concentration-dependent manner (Fig 4C, D). NAD^+^ replenishment also delayed senescence and recovered the levels of histone lactylation in MEFs (Additional file: Fig. S9). By contrast, the compound STF-118804, a selective inhibitor of the NMN producer NAMPT *in vivo*, can promote senescence and diminish histone lactylation (Fig. 4E, F). These findings showed that NAD^+^ content directly regulates histone lactylation abundance.

**Fig. 4.**
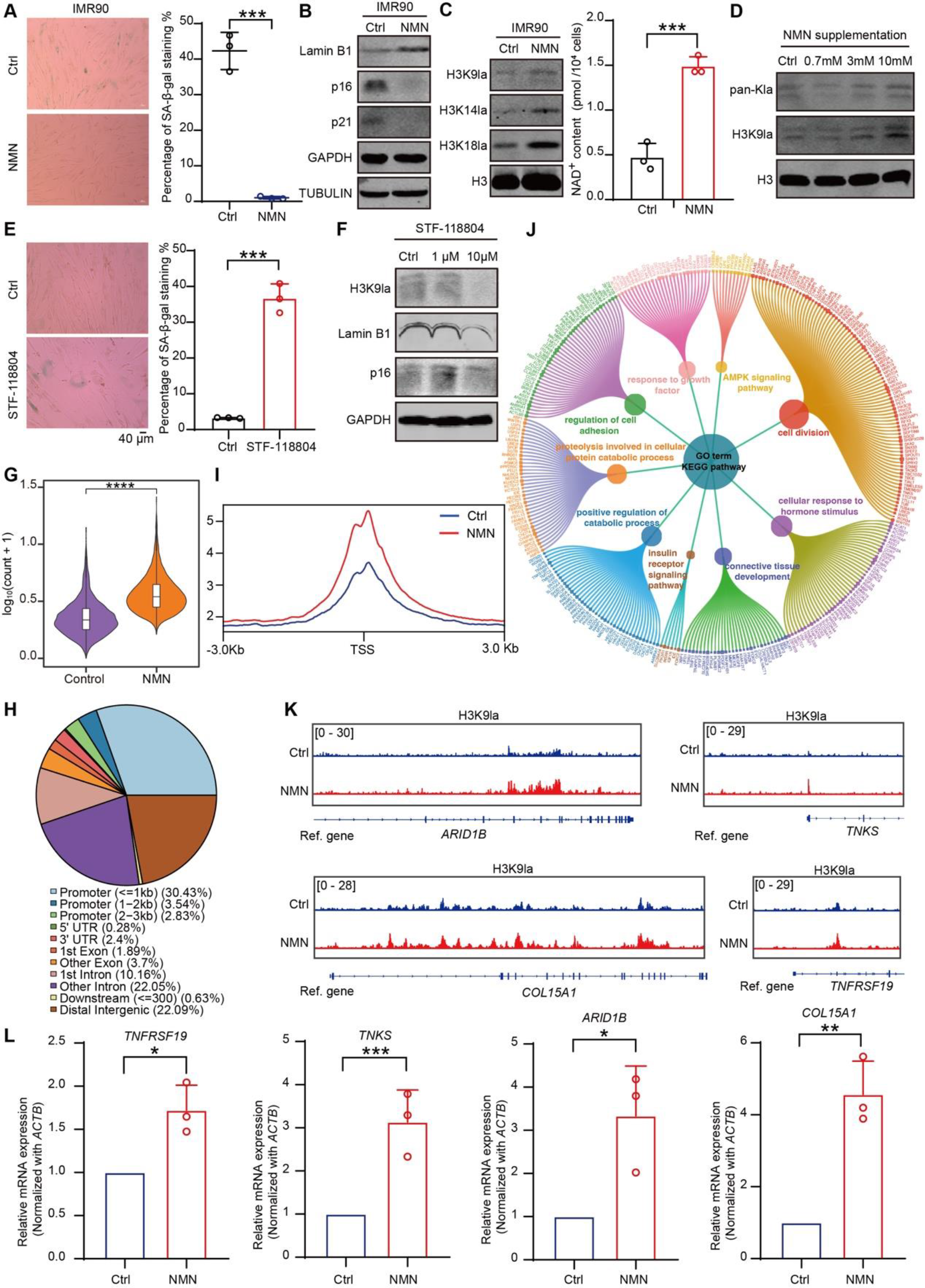
**NAD^+^ content specifically regulates histone lactylation abundance. A**, SA-β-gal staining of IMR90 cells in the presence or absence of NMN. The percentages of SA-β-gal^+^ cells are shown on the right. **B**, Immunoblotting of Lamin B1, p16, and p21 in IMR90 cells in the presence or absence of NMN (10 mM). TUBULIN and GAPDH served as the loading controls. **C**. The left panel shows the immunoblotting of H3K9la, H3K14la, and H3K18la in IMR90 cells in the presence or absence of NMN (10 mM). H3 served as the loading control. The right panel shows the NAD^+^ content in IMR90 cells with or without NMN addition. **D**. Protein levels of total histone lactylation (pan-Kla) and H3K9la in IMR90 cells treated with different concentrations of NMN. H3 served as the loading control. **E**, SA-β-gal staining of IMR90 cells with or without STF-118804 treatment. The percentages of SA-β-gal^+^ cells are shown on the right. **F**. Immunoblotting showing the protein levels of H3K9la, Lamin B1, and p16 in IMR90 cells after STF-118804 treatment. GAPDH served as a loading control. **G**. Violin plot of the signal intensity of the upregulated H3K9la peaks following NMN treatment. ****P < 0.0001. **H**, Distribution of the upregulated H3K9la peaks in genomic elements in IMR90 cells following NMN treatment. **I**, H3K9la signal profiles in the range of 3 kb upstream and 3 kb downstream of TSSs throughout the genome in IMR90 cells in the presence or absence of NMN. **G**, Signal intensity heatmap of the differential H3K9la peaks in IMR90 cells in the presence or absence of NMN. **J**, Functional enrichment analysis of genes near the promoters in which H3K9la was upregulated in IMR90 cells following NMN treatment. The outer circle nodes represent genes related to the differential peaks, and the size of the dot represents the fold change in overall peak signal intensity between IMR90 cells in the presence versus absence of NMN. The size of the dot in the inner circle represents the significance of each pathway, with a larger dot indicating greater significance. **K**, Snapshots of the H3K9la peak signals near *TNFRSF19*, *TNKS*, *COL15A1*, and *ARID1B* in IMR90 cells in the presence or absence of NMN. **L**, RT‒ qPCR of *TNFRSF19*, *TNKS*, *COL15A1*, and *ARID1B* in IMR90 cells in the presence or absence of NMN. The cycle threshold (Ct) values of these genes were normalized to that of *ACTB*. The error bars represent the S.D. of three independent experiments. Two-tailed, unpaired Student’s *t*-tests were performed. \**P* < 0.05, \*\**P* < 0.01, \*\*\**P* < 0.001. TSSs, transcription start sites; NMN, nicotinamide mononucleotide.

Similar to the finding in the young IMR90 cells, there was also a remarkably upward trend in differential H3K9la peaks after NAD^+^ resumption (Fig. 4G). The increased H3K9la signal due to NAD^+^ restoration was mainly distributed in promoters, distal intergenic regions, and introns (Fig. 4H), which was similar to the profiles in replicative senescence and hypoxia (Fig. 1H and Fig. 3A). As expected, the signals of histone lactylation remarkably declined around promoters (Fig. 4I, Additional file: Fig. S10A). The results of pathway enrichment analysis showed that histone lactylation located in promoters after NAD^+^ resumption regulated anti-aging pathways, such as the response to growth factors, cell division, adhesion, and proteolysis (Fig. 4J), several of which were consistent with those uncovered by RNA-seq. The RNA-seq results showed that the genes activated in IMR90 cells following NMN treatment were mainly enriched in pathways involved in the response to growth factors and adhesion. As a comparison, the genes activated upon hypoxia were mainly enriched in the pathways involved in cell cycle and cell division. Additionally, the downregulated genes in senescence were mainly enriched in the cell cycle and the unfolded protein response (Additional file: Fig. S10B). These results demonstrate that NAD^+^-dependent histone lactylation is capable of reprogramming the gene expression profiles of senescent cells to a young state.

Likewise, the promoters of *TNFRSF19*, *TNKS*, *COL15A1*, and *ARID1B* exhibited more H3K9la peaks following NMN treatment, and the mRNA levels of H3K9la-targeted genes were increased (Fig. 4K, L and Additional file: Fig. S10C). Moreover, the NMN-specific responsive enhancers mainly regulated cell morphogenesis, similar to the results obtained from senescence, hypoxia, and 2-DG experiments (Additional file: Fig. S11). These findings confirm that histone lactylation plays an important role in senescence by regulating proliferation and homeostasis, and has disparate regulatory functions in enhancers.

### Running exercise recovers histone lactylation of old skeletal muscle

Given the difference between cellular senescence and the aging of individuals, we further explored the dynamics of histone lactylation in the main metabolic organs of mice during aging. Immunoblotting of histone lactylation in aged organs, including the kidney, liver, heart, and skeletal muscle, verified our hypothesis that the level of histone lactylation declines markedly during aging (Fig. 5A and Additional file: Fig. S12). Correspondingly, H3K14la enrichment was down-regulated in the promoters of the classic histone lactylation-targeted genes *Arg1* and *Eno1* ^15^(Fig. 5B). Similar to the case during cellular senescence, there was no clear correlation between the level of lactate and the level of histone lactylation (Additional file: Fig. S13), suggesting the existence of a regulatory intermediary.

**Fig. 5.**
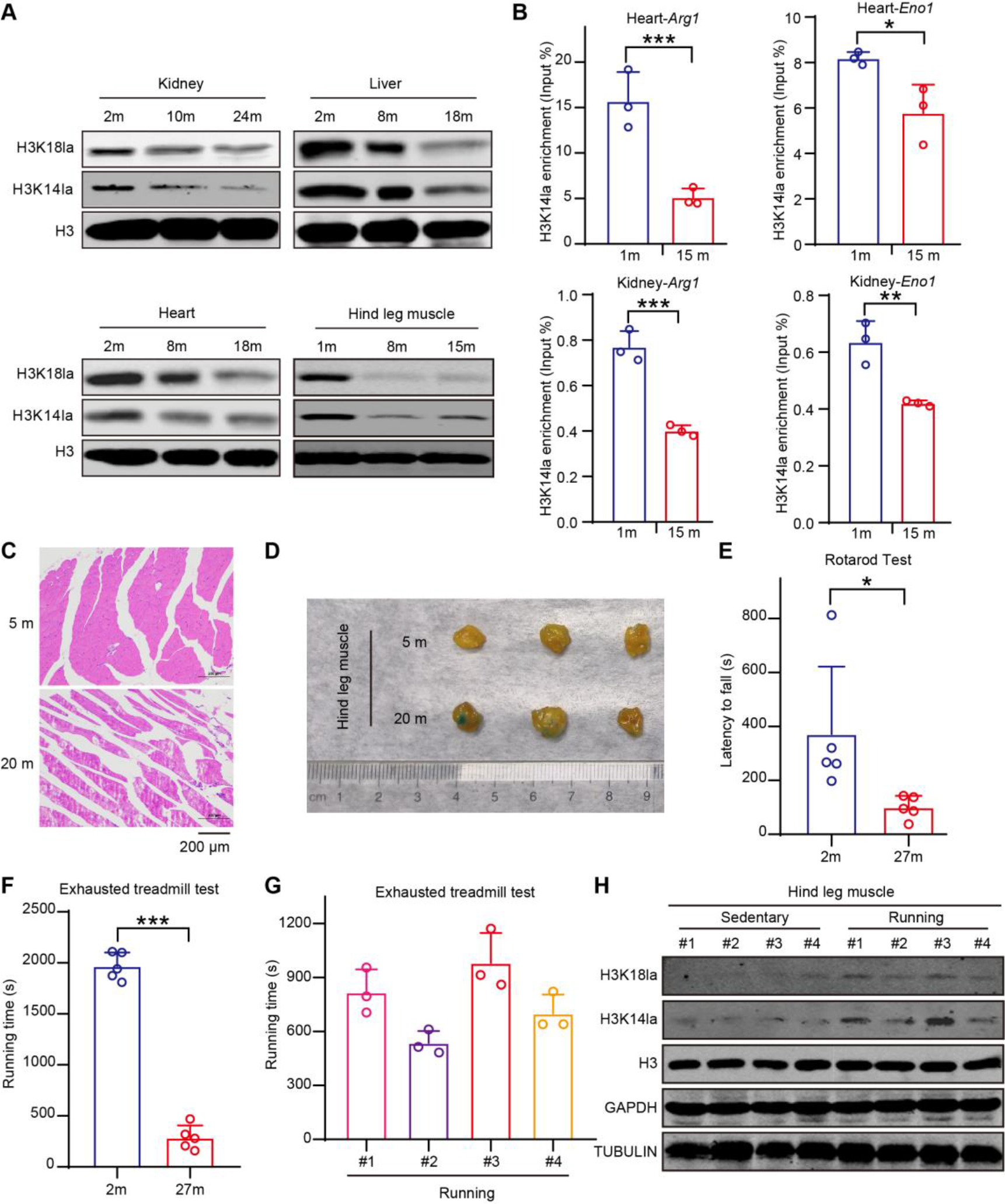
**Exercise recovers global level of histone lactylation. A**, Protein levels of total histone lactylation (pan-Kla), H3K14la, and H3K18la in the kidney, liver, heart, and hindleg muscle of female mice. H3 served as the loading control. **B**, ChIP‒qPCR of the changes in H3K14la signals in the promoters of *Arg1* and *Eno1* during heart and kidney aging. The y-axis represents the H3K14la level measured as the percentage of input (input %). The error bars represent the S.D. of three independent experiments. Two-tailed, unpaired Student’s *t*-tests were performed. \**P* < 0.05, \*\**P* < 0.01, \*\*\**P* < 0.001. **C**, H&E staining of skeletal muscle from the hindleg of 5- and 20-month-old female mice. **D**, SA-β-gal staining of skeletal muscle from the hindleg of 5- and 20-month-old female mice. Three mice were measured in each group. **E**, The rotarod performance test for 2- and 27-month-old male mice. The y-axis represents the persistence time spent on the rotating rod (latency to fall). Five mice were tested in each group. **F**, The treadmill fatigue test for 2- and 27-month-old male mice. The y-axis represents the time taken to fatigue. Five mice were tested in each group. **G**, The treadmill fatigue test for 13-month-old female mice. The y-axis represents the time taken to fatigue. Experimental data were recorded for four mice. The error bars represent the S.D. of three independent experiments performed on different days. **H**, Protein levels of total histone lactylation (pan-Kla), H3K14la, and H3K18la in sedentary and exercised mice. TUBULIN, GAPDH, and H3 served as the loading controls. The four exercised mice were from the treadmill fatigue test in (**G**).

Furthermore, we explored the function of histone lactylation in skeletal muscle aging in terms of the most significant decrease in histone lactylation in this tissue. Skeletal muscle is the primary motor organ; therefore, we hypothesized that histone lactylation likely affects exercise capacity. We evaluated the changes in the structure and motor ability of mouse skeletal muscle during aging and found that fiber fractures accumulated and SA-β-gal^+^ staining increased (Fig. 5C, D); meanwhile, the exercise capacity of aged mice considerably decreased (Fig. 5E, F). Next, we selected middle-aged mice (13 months old) for exercise on a treadmill (Fig. 5G). Interestingly, the histone lactylation level was markedly recovered in mouse skeletal muscle after short-term training (Fig. 5H). Moreover, mice with a higher running ability had higher levels of histone lactylation in skeletal muscle after exercise (Fig. 5G, H). These results indicate that histone lactylation contributes to the function of young skeletal muscle and that a higher abundance of histone lactylation represents better exercise performance in mice.

### Histone lactylation accumulated in exercised skeletal muscle reconstructs cell landscape

Next, to substantiate that histone lactylation is important for the youthful appearance of skeletal muscle, we performed single nuclei RNA sequencing (snRNA-seq) using young and old skeletal muscle tissues to analyze the changes in the cell landscape during skeletal muscle aging. We acquired 7,141 cells that were qualified by gene numbers (Additional file: Fig. S14A) and analyzed the cell composition of skeletal muscle. A total of 10 cell types were verified: type IIa myonuclei identified by the expression of *Myh2* and *Ppara*, type IIb myonuclei identified by the expression of *Mylk4* and *Mical2*, type IIx myonuclei identified by the expression of *Myh1*, unidentified myonuclei identified by the expression of *Sh3d19*^39^, fibro-adipogenic progenitors identified by the expression of *Abca8a*, slow myonuclei identified by the expression of *Myh7*, myotendinous junction identified by the expression of *Col22a1*, neuromuscular junction identified by the expression of *Colq* and *Ano4*, adipocytes identified by the expression of *Pde3b* and *Acsl1*, and endothelial cells identified by the expression of *Ptprb* (Fig. 6A, B, and Additional file: Fig. S14B).

**Fig. 6.**
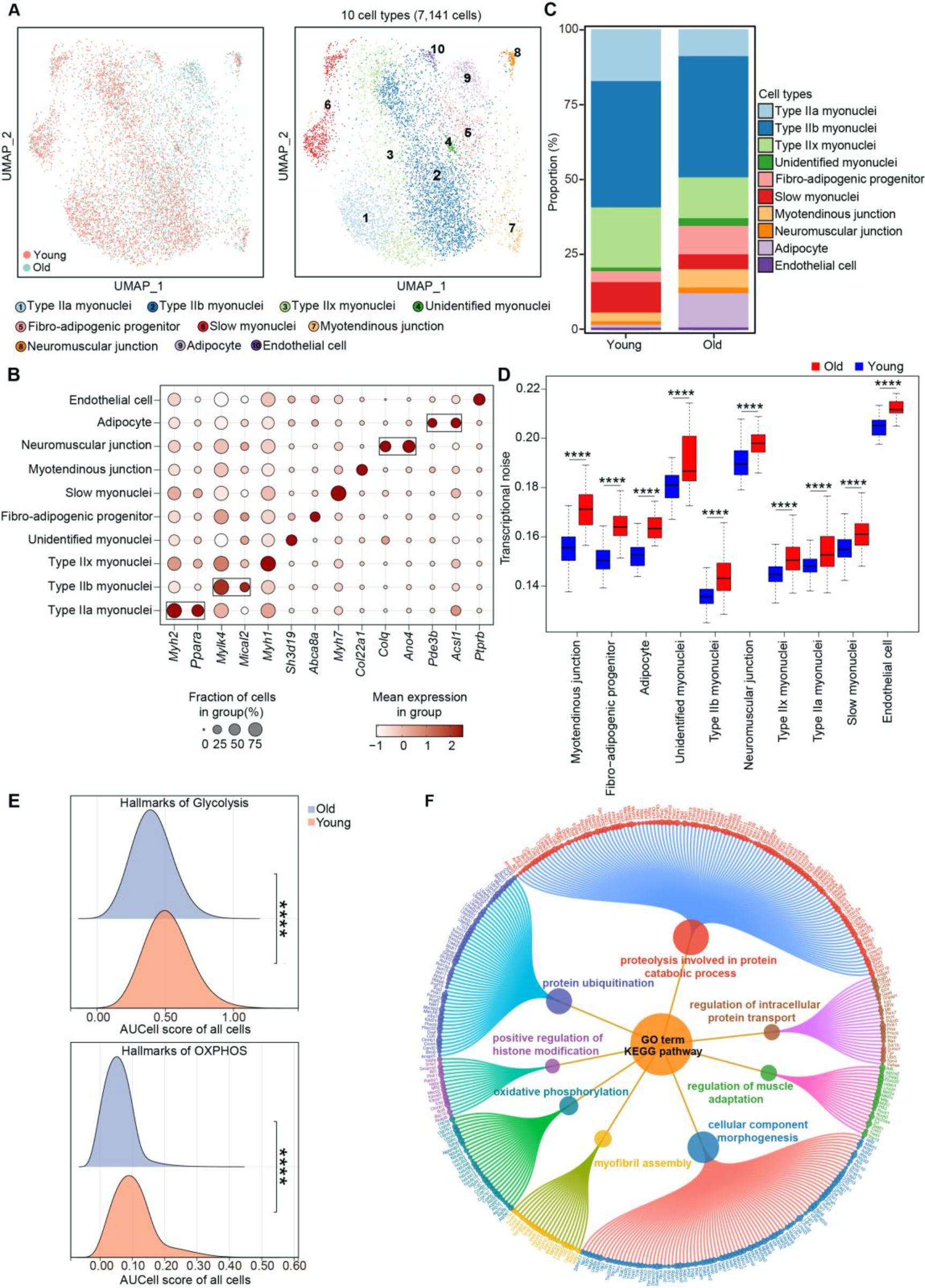
**Cell activities undergo massive degeneration during skeletal muscle aging. A**, UMAP visualization of the 10 major cell types from the snRNA-seq data for young and old female mice. The left part shows the UMAP plot of the sample origin. **B**, Dot plots of marker gene expression in the 10 major cell types in (**A**). The x-axis represents the marker gene, and the y-axis represents the cell type. **C**, Stacked bar chart of the cell ratios of young and old female mice. **D**, Box diagram showing the levels of change of transcriptional noise in all kinds of identified cell types along with skeletal muscle aging. **E**. Ridge plot showing the enrichment score for the glycolysis and OXPHOS hallmark genes in the skeletal muscle of young and old samples. OXPHOS, oxidative phosphorylation. **F**, Functional enrichment analysis of the downregulated genes in the slow myonuclei of young versus old mice. The outer circle nodes represent the DEGs, and the dot size represents the fold change in gene expression. The dot size in the inner circle represents the significance of the enriched pathway, with a larger dot indicating greater significance.

Thereinto, the proportions of type IIa myonuclei, type IIx myonuclei, and slow myonuclei had obvious downward trends. By contrast, the proportions of unidentified myonuclei, fibro-adipogenic progenitors, myotendinous junction, adipocytes, and endothelial cells were increased (Fig. 6C). These changes in cell proportions were consistent with the macroscopic transitions of skeletal muscle aging, namely, muscle mass loss and functional decline while gaining fat. We also found that the transcriptional noise^40^ of various cell types increased significantly during skeletal muscle aging (Fig. 6D), indicating the augmentation of transcription disorder and disturbance of gene expression. Furthermore, the metabolic capacity of old skeletal muscle was significantly decreased, as revealed by decreased expression of hallmark genes involved in glycolysis and OXPHOS (Fig. 6E, Additional file: Fig. S15).

In view of the common downward trend between the proportion of cells promoting muscle function and the abundance of histone lactylation in aging, we hypothesized that there is a potential causal relationship between the two. Subsequently, we focused on the slow myonuclei and performed further functional analysis due to the superior metabolic capacity of these myonuclei. Along with aging, the overall activity of gene expression decreased in slow myonuclei (Additional file: Fig. S14C), especially the genes for proteolysis, protein transport, muscle adaptation, myofibril assembly and component morphogenesis (Fig. 6F).

To investigate the underlying mechanism of histone lactylation in regulating skeletal muscle function, we trained middle-aged mice using long-term running exercise. Compared with that in the untrained controls, the percentage of SA-β-gal^+^ cells in skeletal muscle was reduced in the trained animals (Fig. 7A). Subsequently, single-nuclei RNA sequencing was performed to evaluate the potential function of histone lactylation in skeletal muscle^39^. Sequencing data showed reliable quality features following analysis according to gene number, unique molecular identifier (UMI) number, and percentage of mitochondrial genes (Additional file: Fig. S16A). After computational quality control, 8,273 cells were remained for further analyses. A total of 12 cell types were discovered in the skeletal muscle samples from sedentary and exercised mice using canonical markers and highly expressed specific genes: type IIa myonuclei, type IIb myonuclei, type IIx myonuclei, unidentified myonuclei, slow myonuclei, myotendinous junction, neuromuscular junction, adipocytes, endothelial cells, fibro-adipogenic progenitors, satellite cells, and smooth muscle cells (Fig. 7B). Specific markers for the corresponding cell types had preferential expression patterns (Fig. 7C and Additional file: Fig. S16B). In addition, the levels of hallmark biological processes that contribute to skeletal muscle regeneration, e.g., glycolysis, oxidative phosphorylation (OXPHOS), and hypoxia, were considerably increased, with a particularly remarkable phenomenon observed in slow myonuclei (Fig. 7D and Additional file: Fig. S17).

**Fig. 7.**
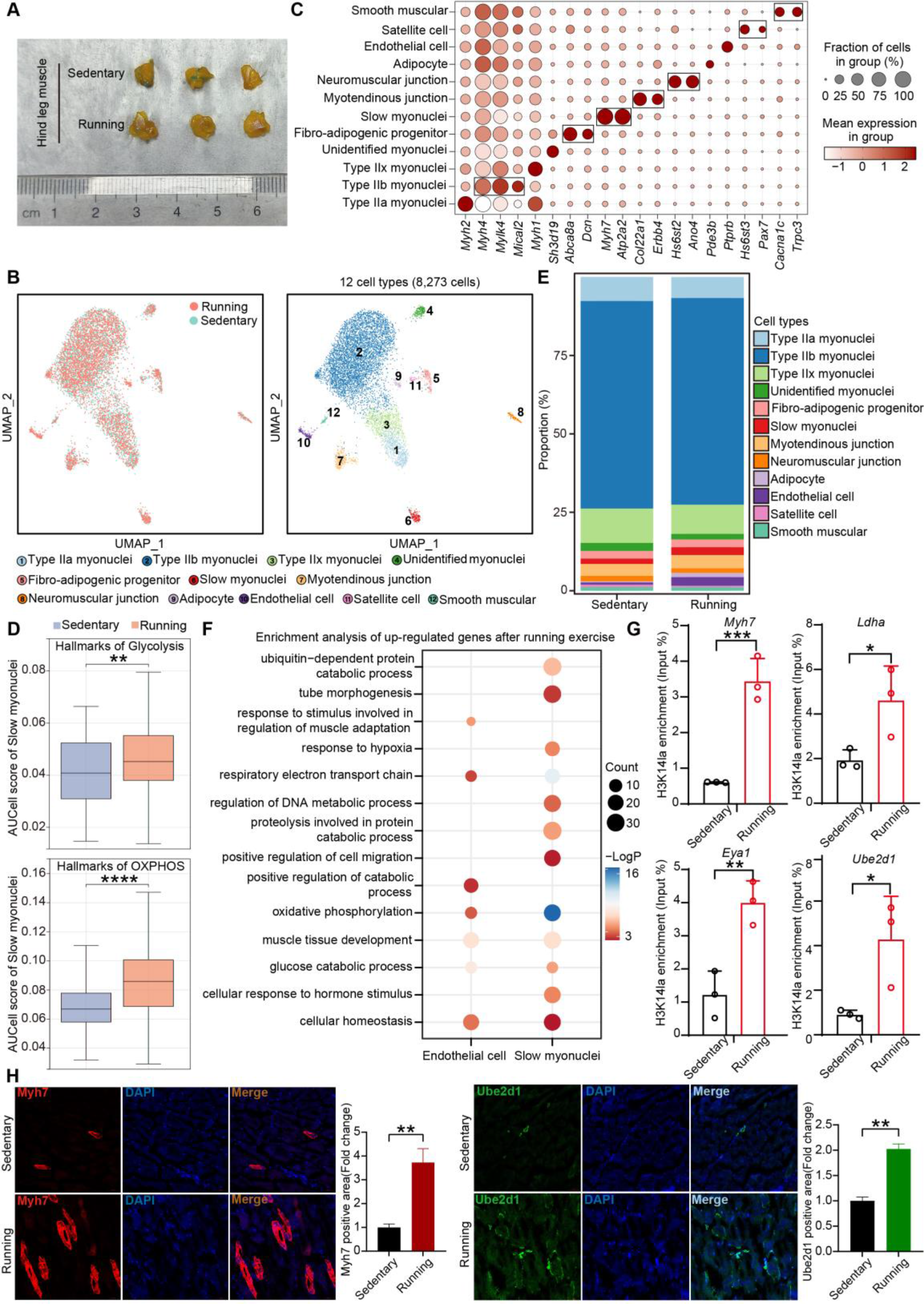
**Exercise facilitates histone lactylation resumption and cell landscape reconstruction. A**, SA-β-gal staining of skeletal muscle from the hindlegs of sedentary and exercised female mice. Three mice were measured in each group. **B**, UMAP visualization of the 12 major cell types from the snRNA-seq data for sedentary and exercised female mice. The left part shows the UMAP plot of the sample origin. **C**, Dot plots of marker gene expression in the 12 major cell types in (**B**). The x-axis represents the marker gene, and the y-axis represents the cell type. **D**, Box plot showing the expression of hallmark glycolysis and OXPHOS genes in the skeletal muscle of sedentary and running female mice. OXPHOS, oxidative phosphorylation. **E**, Stacked bar chart of the cell ratios of sedentary and exercised female mice. **F**, Functional enrichment analysis of the upregulated genes in endothelial cells or slow myonuclei after exercise. **G**, ChIP‒qPCR of H3K14la enrichment at the promoters of *Myh7*, *Ldha*, *Eya1*, and *Ube2d1* in sedentary and exercised mice. The y-axis represents the H3K14la signal measured as the percentage of input (input %). **H**, Immunofluorescence of Myh7 and Ube2d1 in skeletal muscle from sedentary and running mice. The error bars represent the S.D. of three independent experiments. Two-tailed, unpaired Student’s *t*-tests were performed. \**P* < 0.05, \*\**P* < 0.01. snRNA-seq, single nuclei RNA-seq.

Significant rearrangement of cell composition occurred in the skeletal muscle from running mice in comparison with that from sedentary mice. The proportions of slow myonuclei and endothelial cells were markedly increased after exercise training (Fig. 7E), demonstrating that exercised mice acquired young muscle functions such as strength, endurance, and myofiber regeneration. Next, we analyzed the enriched pathways related to the upregulated genes in these three cell types, which were found to be involved in the regulation of superior muscle function^41^, such as energy metabolism, proteolysis, and the response to hormone stimuli (Fig. 7F). To validate the connection between histone lactylation and skeletal muscle function, we evaluated H3K14la enrichment in promoters that regulate the upregulated genes in these cells. Accordingly, the level of H3K14la was increased in the promoters of *Myh7*, *Ldha*, *Eya1*, and *Ube2d1* (Fig. 7G). The protein expression of Myh7 and Ube2d1 increased correspondingly (Fig. 7H), in contrast to their downregulation during aging (Additional file: Fig. S18). These findings suggest that histone lactylation plays a positive role in the vigorous functions of active skeletal muscle by reconstructing cell composition during running exercise.

### Long-term running rewires cell communication in skeletal muscle

Next, considering the remarkable reconstruction of the cell landscape mediated by histone lactylation after running exercise, we wondered whether cell communication was rewired under this circumstance. Surprisingly, we found that the interaction network among skeletal muscle cells was rearranged in running-exercised mice. Particularly noteworthy were the observed increases in interactions of slow myonuclei, myotendinous junction, fibro-adipogenic progenitors, satellite cells, and type IIb myonuclei with other cell types. These multifaceted and intricate interactions likely play a vital modulatory role in muscle development and regeneration. Simultaneously, reductions in the interactions of adipocytes and endothelial cells with other cell types were observed (Fig. 8A). The results of the interaction strength analysis were consistent with the corresponding numbers (Additional file: Fig. S19A). Overall, the interaction numbers and strength in skeletal muscle cells were dramatically improved in running mice compared with sedentary mice (Fig. 8B, Additional file: Fig. S20A). By contrast, both the interaction numbers and interaction strength were decreased in old skeletal muscle (Fig. 8B, Additional file: Fig. S19B, Fig. S20B). Moreover, most of differential expression genes were up-regulated in almost all cell types, which may suggests that running exercise exerts a profound impact on the systemic metabolism and physiological performance of skeletal muscle (Fig. 8C).

**Fig. 8.**
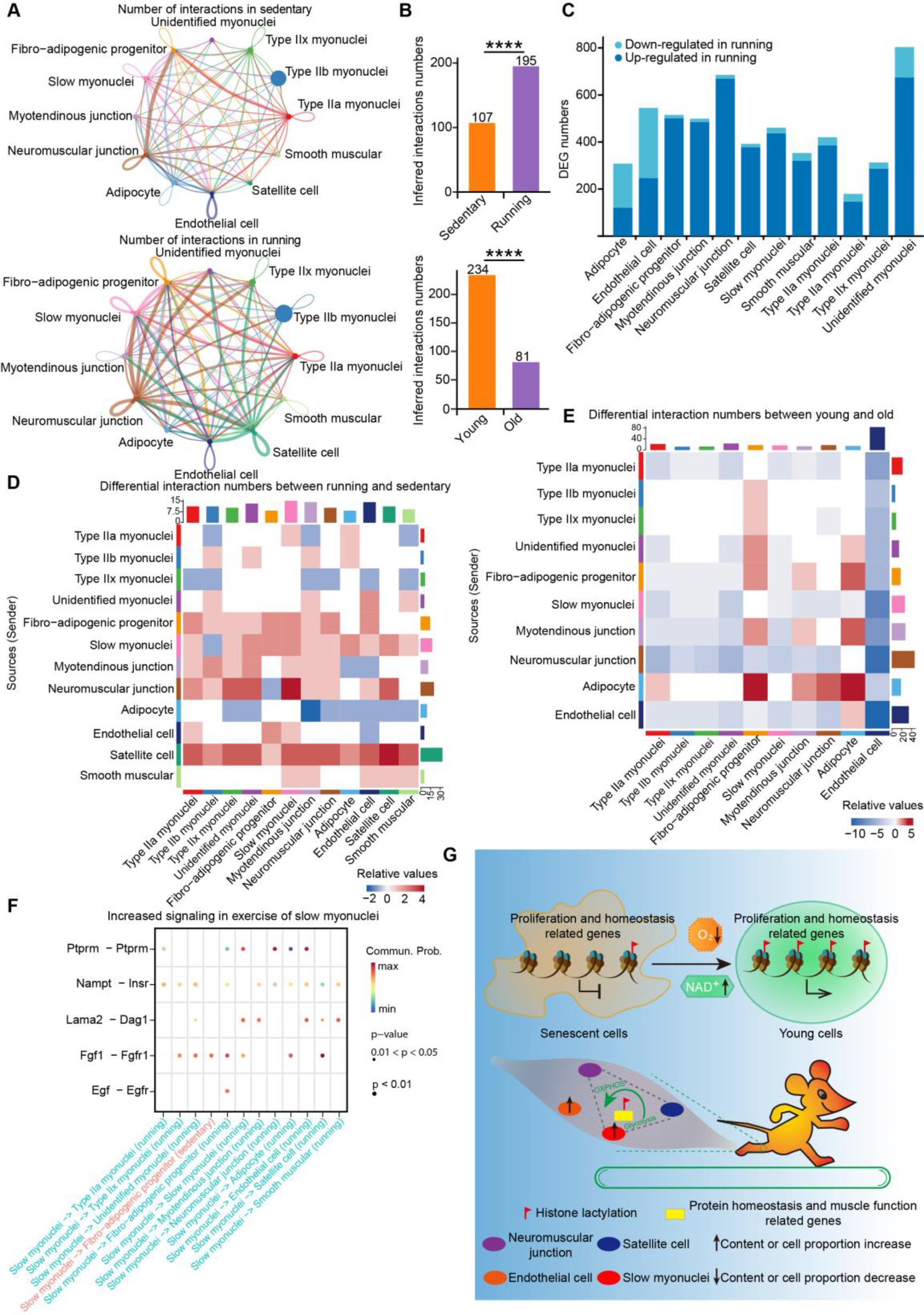
**Cell communications are enhanced in exercised skeletal muscle. A**, Analysis of interaction numbers in skeletal muscle cells of sedentary (upper) and running (lower) mice using snRNA-seq data. Each dot represents the cell type, and the thickness of the line represents the interaction number. **B**, Bar graph showing the changes in cell interaction numbers during exercise or aging in female mice. The Wilcoxon rank sum test was performed, \*\*\*\**P* < 0.0001. **C**, Statistics for the numbers of down- or upregulated DEGs in different cell types after running exercise. **D**, Heatmap showing the numbers of differential interactions between cell types in sedentary and running samples. Red (blue) indicates that there are more (fewer) cellular interactions in the running samples than in the sedentary samples. The color shade represents the difference in interaction number between cell types. **E**. Heatmap showing the numbers of differential interactions between cell types in young and old samples. Red (blue) indicates that there are more (fewer) cellular interactions in the old samples than in the young samples. The color shade represents the difference in interaction numbers between cell types. **F**, Dot plot showing the differentially upregulated ligand‒receptor interactions in slow myonuclei of running samples. **G**. Hypothetical working model showing the anti-aging role of histone lactylation. In replicative senescence, the abundance of histone lactylation decreases obviously near anti-aging genes, including proliferation- and homeostasis-related genes. Restoration of histone lactylation by physiological hypoxia (3% O_2_) or NAD^+^ resumption can delay senescence by reprogramming the gene expression profile. In skeletal muscle aging, running exercise recovers the lost histone lactylation, which reactivates protein homeostasis and muscle function-related genes, contributing to reconstruction of the cell landscape and communication. Inside, the interactions of neuromuscular junction cells with slow myonuclei and satellite cells are strengthened, and the proportions of slow myonuclei and endothelial cells evidently increase. Partial vector elements for drawing were downloaded from the Reactome Pathway Knowledgebase^55^.

When further analyzing the cell-cell communication based on differentially expressed gene, we found that the interactions of neuromuscular junction with nearly all other cell types were strengthened, except those with adipocytes and fibro-adipogenic progenitors. Specifically, the interactions among neuromuscular junction, slow myonuclei, and satellite cells were the most prominent. The interactions of slow myonuclei with endothelial cells and myotendinous junction were mildly enhanced (Fig 8D, Additional file: Fig. S20C). However, the cell communication activity was quite low in old skeletal muscle among these cells (Fig. 8E, Additional file: Fig. S20D), indicating that exercise reactivates the communication networks to withstand the aging program. Regarding communication signaling, increased Ptprm-Ptprm, Nampt-Insr, Lama2-Dag1, Fgf1-Fgfr2, and Egf-Egfr signaling mediated the interactions of slow myonuclei with other cell types, including neuromuscular junction and endothelial cells (Fig. 8F). Furthermore, the Nampt-Insr interaction can promote the biosynthesis of NAD^+^, exert anti-inflammatory effects, and play a crucial role in promoting cell metabolism^42^. Fgf1-Fgfr1 also contributes to promoting cell proliferation, differentiation^43^, and the formation of new blood vessels^44^. And the increased signaling of Ptprm-Ptprm and Egf-Egfr mediated the interactions of endothelial cells with type IIa myonuclei, fibro-adipogenic progenitors, and slow myonuclei (Additional file: Fig. S21). These findings demonstrate that cell-cell communications are rewired after running exercise due to re-engineering of the cell landscape via histone lactylation. The reconstruction of cell composition and communication promote a youthful state of skeletal muscle.

## DISCUSSION

Our discoveries reveal the characteristics and function of histone lactylation in cellular senescence and skeletal muscle aging. Herein, we report a great decline in histone lactylation during senescence and aging. Physiological hypoxia and NAD^+^ resumption- led metabolism reprogramming for primary cells manage to recover the histone lactylation level and revealed a positive correlation of this novel histone modification with a youthful cellular state. The genome-wide profiles of histone lactylation show that the anti-aging function of histone lactylation is implemented via activation of pathways related to proliferation and homeostasis. More importantly, the outcomes of running exercise illustrate that youthful function can be regulated by histone lactylation in mouse skeletal muscle (Fig. 8G).

Histone lactylation participates in senescence by regulating cellular homeostasis, proliferation, and interaction with the environment through genes involved in proteolysis, DNA repair, cell cycle, and the response to growth factors (Fig. 2F, Fig. 3I, and Fig. 4H). Several functions of histone lactylation, including the promotion of homeostasis in M1 macrophages^15^ and the activation of proliferation in ocular melanoma cells, have also been reported^20^. In functional genomic regions, histone lactylation not only regulates promoters but also influences the activity of enhancers, a phenomenon that has been reported in other cell lines^27^. Our analyses reflect the specificity of enhancers for different environmental alterations regulated by histone lactylation. The regulatory function of histone lactylation on chromatin structure is also worthy of exploration. Our discoveries in individuals uncover new functions of histone lactylation in the regulation of skeletal muscle regeneration and differentiation (Fig. 7 and Additional file: Fig. S20). Moreover, it will be of great interest to investigate the role of histone lactylation in other organs given its widespread distribution (Fig. 1G–I and Fig. 6A).

Our findings demonstrate that the abundance of histone lactylation markedly decreases in MEFs, HUVECs, and IMR90 cells during senescence, as well as in mouse heart, liver, kidney, and skeletal muscle tissues during aging, reflecting the conservation of this modification across different cell types and species. However, we cannot rule out an inconsistent tendency of histone lactylation levels that depends on the types of cells, organs, and senescence or at other histone sites that are not included in this study. The relationships between histone lactylation and several modifications have been described in other models^19, 21, 27^, suggesting that the coordination of histone lactylation with other epigenetic modifications, such as histone acetylation and DNA methylation, also deserves investigation in the contests of senescence and aging. Our results and those previously reported by others indicate that histone lactylation functions in a plethora of physiological and pathological processes^15–22, 24, 27, 45, 46^.

It should be noted that changes in the endogenous lactate level and histone lactylation abundance did not correlate with each other during senescence and aging (Fig. 1G–I, Fig. 6A, and Additional file: Fig. S2A, Fig. S18). Interestingly, it has been demonstrated that glycolysis is uncontrolled in senescent cells and that the lactate level may not represent the actual glycolytic flux^47, 48^. Further investigation is warranted to unveil the myths underlying lactate-related metabolism and lactylation since lactate is an indirect substrate for histone lactylation due to its multiple uses, e.g., as energy, a signaling molecule, and a source of lactylation^15, 49^. Instead, the lactyl group for histone lactylation is directly provided by L-lactyl-CoA in a p300-dependent manner^15^. Moreover, we have shown that the histone lactylation level is dependent on NAD^+^ concentration (Fig. 4B), implying that NAD^+^ could be a potential factor of metabolic intermediates that can regulate histone lactylation in senescence and other life processes.

The snRNA-seq analyses revealed that the increased histone lactylation induced by exercise reverses the adverse changes in cell composition and communication networks during skeletal muscle aging. Histone lactylation can delay aging by activating genes involved in protein homeostasis pathways, which is similar to the mechanism by which senescence is antagonized. Furthermore, the biosynthesis of NAD^+^ may be improved in exercised skeletal muscle as manifested by enhancement of the Nampt-Insr interaction. This interesting clue is like the result of NMN treatment in cells. Despite the known importance of elucidating the molecular mechanisms by profiling the genome-wide enrichment of histone lactylation in skeletal muscle aging and running exercise models, we are hampered by the poor efficiency of the commercially available antibody for skeletal muscle.

In summary, our data expand the body of knowledge regarding the roles of histone lactylation in senescence and aging regulation and provide the possibility to not only delay the progression of senescence but also alleviate aging phenotypes in tissues and facilitate functional improvements of skeletal muscle via metabolism manipulation.

## Methods

### Cell culture

IMR90 cells were purchased from the American Type Culture Collection (ATCC), HUVECs were purchased from AllCells, and MEF cells were isolated from 12.5–14- day-old embryos of C57BL/6 mice. IMR90 and MEF cells were cultured in DMEM (Fisher Scientific) supplemented with 10% fetal bovine serum (FBS, Fisher Scientific), and nonessential amino acids (NEAA, 100×, Macgene) were added for IMR90 cell culture. For physiological hypoxia experiments, IMR90 and MEF cells were cultured in an incubated hypoxic chamber (STEMCELL, 27310) containing mixed gas (3% O2, 5% CO2, 92% N2). All cells were cultured at 37°C with 5% CO 2.

### Mouse models

C57BL/6 mice were purchased from Charles River. All animal experiments met the requirements of the Institutional Animal Care and Use Committee of Peking University. For the rotarod performance test, the initial speed was set to 5 rpm, and the final speed was 32 rpm, with an acceleration time of 300 s. Five mice in each group were tested on three consecutive days, and the latency to fall (s) was recorded. For the treadmill fatigue test, four mice were tested on three separate days, with an interval of one day. The slope of the treadmill was set to 5°, and the speed (m/min) was increased as follows: 10 (2 min), 14 (5 min), 16 (5 min), 18 (5 min), and 20 (5 min). The running time to fatigue was recorded. For long-term running exercise using a treadmill, the speed was set to 9 m/min, and the running time was 30 min for each training session. Running exercise lasted for 2 months, with 2–3 days of rest after each training session. Five exercised mice ran for 2 min at 6 m/min to adapt to the treadmill each time.

### Senescence-associated β-galactosidase staining (SA-β-gal staining)

Cells were seeded onto 6-well plates and cultured to the required density, after which SA-β-gal staining was performed using a Senescence Cells Histochemical Staining Kit (Sigma, CS0030). Cells were incubated with the staining mixture overnight at 37°C. Images were acquired using a DMI 6000B microscope with 10 × 10 magnification (Leica). The Image J software (NIH) was used to analyze the percentage of SA-β-gal^+^ signals. The experimental procedures for SA-β-gal staining of skeletal muscle were performed in 15-mL tubes, and images were acquired using a camera.

### Immunofluorescence

Cells were seeded onto sterile coverslips in 10-cm plates and cultured to the required density. The cells were fixed for 10 min with 4% paraformaldehyde, blocked for 30 min with 1% bovine serum albumin (BSA) at room temperature, and incubated overnight at 4°C with an anti -Ki67 antibody (Abcam, ab15580, 1:500). The next day, the cells were incubated with Alexa Fluor 594 Donkey anti-Rabbit IgG (Life Technologies) for 2 h at room temperature and then stained with 1 ng/μL DAPI for 3 min. Immunofluorescence images were acquired from sealed coverslips using a fluorescent microscope (Leica).

### Immunoblotting

TRIzol® (Invitrogen, 15596018) was used to extract proteins from cells or tissues, which were isolated by SDS-PAGE and transferred to nitrocellulose membranes. The membranes were incubated with 5% skimmed milk for 1 h, followed by specific antibodies overnight at 4°C: anti -Lamin B1 (Proteintech, 12987-1-AP, 1:1000), anti- p16 (Proteintech, 10883-1-AP, 1:500), anti-p21 (Santa Cruz Biotechnology, sc397, 1:500), anti-β-ACTIN (Santa Cruz Biotechnology, sc-47778, 1:1000), anti-α-TUBULIN (Sigma, T6199, 1:5000), anti-GAPDH (Proteintech, 60004-1-Ig, 1:10000), anti-L-lactyllysine (pan-Kla, PTM BIOLABS, PTM1401, 1:1000), anti-H3K9la (PTM BIOLABS, PTM1419RM, 1:1000), anti-H3K14la (PTM BIOLABS, PTM1414, 1:1000), anti-H3K18la (PTM BIOLABS, PTM1406, 1:1000), and anti-H4K12la (PTM BIOLABS, 1:1000, PTM1411RM). The next day, the membranes were incubated with IRDye 800CW goat/donkey anti-mouse/rabbit antibodies (LI-COR Biosciences, 926- 32210, 1:10000) for 2 h at room temperature. Images were acquired using an Odyssey Infrared Imaging System (LI-COR).

### CUT&Tag sequencing (CUT&Tag-seq)

Cells were harvested by trypsin digestion at room temperature and then counted. CUT&Tag experiments were performed according to the CUT&Tag Assay Kit manual (Vazyme, TD903). Briefly, cell pellets were collected at 600 × g for 5 min, washed with wash buffer, and incubated with ConA beads for 10 min at room temperature in 8-strip tubes. Subsequently, the cell–ConA bead complexes were incubated with anti- H3K9la (PTM BIOLABS, PTM1419RM) at 4°C overnight and with a secondary antibody for 1 h at room temperature. The pA/G-Tnp transposon mediated the transposition reaction, and TTBL mediated DNA fragmentation. DNA extraction beads were used to extract CUT&Tag-enriched DNA, which was then subjected to library amplification for Illumina sequencing. The PCR products were purified using VAHTS® DNA Clean Beads (Vazyme, N411) and sequenced on the NovaSeq 6000 PE150 platform at GENEWIZ.

### RNA-seq

For bulk RNA-seq, total RNA was extracted from cells using TRIzol® (Invitrogen, 15596018). The cDNA library construction and Illumina sequencing were performed by Novogene (HiSeq PE150) or GENEWIZ (NovaSeq 6000 PE150). For single-nuclei RNA sequencing (snRNA-seq), fresh mouse skeletal muscle was frozen in liquid nitrogen and used for library construction and sequencing by BGI Genomics. Five mice in each group were used for snRNA-seq.

### Chromatin immunoprecipitation (ChIP)

Approximately 0.04 g of fresh or frozen tissue was minced using a blade and collected into a 1.5-mL DNA LoBind® Tube (Eppendorf), to which 1 mL of PBS was added.

Crosslinking was performed with 1% formaldehyde for 5 min, after which the reaction was terminated by incubation with 0.125 M glycine for a further 5 min. Crosslinked tissue fragments were washed twice with PBS, and nuclear lysis buffer (50 mM Tris-Cl, 10 mM EDTA, 1% SDS) supplemented with protease inhibitor cocktail was added. The tissue was homogenized with an electric grinder and sonicated using a Bioruptor® (Diagenode). The released DNA fragments were diluted in IP dilution buffer (20 mM Tris-Cl, 2 mM EDTA, 150 mM NaCl, 1% Triton-X100) supplemented with protease inhibitor cocktail and incubated with anti-H3K14la (Abcam, PTM1414). Dynabeads™ Protein G (Invitrogen, 10001D) were used to pull down the chromatin–antibody complexes, and immunoprecipitated DNA was isolated using the phenol‒chloroform extraction method.

### qPCR

All-in-One Supermix (TransGen Biotech, AT341) was used to generate cDNA from total RNA. cDNA or ChIP DNA was measured by qPCR using a LightCycler® 96 Instrument (Roche) after the addition of SYBR Green premix (Vazyme, Q712). The primers used for RT‒qPCR and ChIP‒qPCR are listed in Tables S1 and S2, respectively.

### Hematoxylin and eosin (H&E) staining

Skeletal muscle was paraffin embedded, sectioned, and stained sequentially with hematoxylin and eosin. Images were acquired using a NIKON Eclipse Ci microscope (Nikon).

### Data analysis Bulk RNA-seq

The raw sequencing data were processed using trim_galore (version 0.6.7) to remove adapters and low-quality reads with the following parameters: -q 25 --phred33 -- stringency 4 --length 36 -e 0.1. The trimmed reads were then mapped to the hg38 or mm10 reference genome using hisat2 (version 2.2.1), and the SAM files were converted to sorted BAM files using samtools (version 1.7). The gene expression count matrix was obtained using featureCounts (version 2.0.1) and normalized with DESeq2 (version 1.34.0). The thresholds of differentially expressed genes were set to a log2(fold change) > 0.5 and an adjusted *P* value < 0.001 for 2-DG treatment, while the thresholds for the other treatments were set to a log2(fold change) > 0.5 and a *P* value < 0.05. The volcano plot of differentially expressed genes was generated using ggplot2 (version 3.3.5). The R package GSVA (version 1.34.0) was used to evaluate the pathway enrichment of gene sets related to aging, which were downloaded from the Molecular Signatures Database (MSigDB, version 3.0).

### CUT&Tag-seq

Clean data was mapped to the hg38 reference genome using bowtie2 (version 2.2.5) with the following parameters: --end-to-end --very-sensitive --no-mixed --no-discordant --phred33 -I 10 -X 700. The sorted BAM files were converted to bigWig format using bamCoverage (version 3.5.1) with the following parameters: -- normalizeUsing RPKM. The normalized signal was visualized in Integrative Genomics Viewer (IGV, version 2.11.3). In addition, deepTools (version 3.5.1) was used to calculate signal scores ±3 kb from the transcriptional start site. Peak calling was performed using MACS2 (version 2.2.7.1) with the parameters -f BAMPE -g hs -q 0.05. The DESeq2 method in DiffBind (version 2.14.0) was used to identify differential peaks as defined by a log2(fold change) > 0.5 and a *P* value < 0.05. Peak distribution and annotation were analyzed using the R package ChIPseeker (version 1.22.1). Human enhancer annotations were obtained from the ENCODE database, and the peaks located in enhancers were identified using bedtools (version 2.30.0).

### SnRNA-seq

The FASTQ-format data were aligned to the mm10 reference genome and annotated using STAR (2.7.1). The filtered gene expression count matrix was obtained for downstream analysis using PISA (version 0.12)^50^. For each dependent sample, genes detected in at least 3 cells expressing at least 200 genes were kept. Subsequently, cells with the following three criteria were retained: more than 200 detected genes, fewer than 6000 detected genes, and fewer than 6% mitochondrial genes.

The RunHarmony function in the R package harmony (version 0.1.0) was used to remove batch variances with the parameter max.iter.harmony = 20. Meanwhile, mitochondrial genes and ribosomal genes were removed to avoid noise in downstream analysis. The first 30 principal components (PCs) were selected to perform dimensionality reduction, and then the nearest-neighbor graph was constructed using the FindNeighbors function. Cell clusters were identified using the FindClusters function, and the UMAP plot was generated using the RunUMAP function and visualized with the DimPlot function. The DEGs with a *P* value < 0.05 for each cluster were identified with the FindAllMarkers function using the Wilcoxon rank sum test (min.pct = 0.1, logfc.threshold = 0.25). Cell types were annotated using canonical marker genes^39, 51, 52^ including *Myh2* for type IIa myonuclei; *Myh4*, *Mylk4*, and *Mical2* for type IIb myonuclei; *Myh1* for type IIx myonuclei; *Col22a1*, *Ankrd1*, and *Erbb4* for myotendinous junction; *Dcn*, *Abca8a*, and *Scara5* for fibro-adipogenic progenitors; *Sh3d19* for unidentified myonuclei; *Myh7* and *Atp2a2* for slow myonuclei; *Pecam1* and *Ptprb* for endothelial cells; *Hs6st2*, *Ano4*, *Colq*, and *Chrne* for neuromuscular junction; *Pde3b* and *Acsl1* for adipocytes; *Cacna1c* and *Trpc3* for smooth muscle cells; and *Hs6st3* and *Pax7* for satellite cells. We used AUCell in the R package irGSEA (version 1.1.3) to score the activity of hallmark gene sets in MSigDB. The cell-cell communication inference and analysis of snRNA-seq were performed with CellChat (version 1.0.0)^53^. We performed the standard workflow to predict interactions between multiple cell types by using the mouse secretion signaling database. In addition, the differences in the number and intensity of interactions between samples were performed by Wilcoxon rank sum test.

### Functional enrichment analysis

Metascape^54^ was used to perform gene function enrichment analysis. GO Biological Processes analysis and KEGG enrichment analysis were performed with the parameter *P* value cutoff = 0.01.

## Data availability

All sequencing data from this study are available under the accession number GSE226008.

## Acknowledgments

We thank Prof. Yifu Qiu, Dr. Jingfei Yao, and Shao Zhou for assistance with the treadmill tests. We are grateful to Dr. Peng Du and Xuehui Lyu for help with the CUT&Tag experiments. We appreciate Prof. Chenjian Li, Dr. Yuanyi Dai, and Dr. Yuanxiu Liu for technical support in the rotarod test. Our work was supported by the National Key Research and Development Project (Grant No. 2021YFA0909300) and the National Natural Science Foundation of China (Grant No. 32241006).

## References

1 Kumari, R. & Jat, P. Mechanisms of Cellular Senescence: Cell Cycle Arrest and Senescence Associated Secretory Phenotype. Front Cell Dev Biol 9, 645593, doi:10.3389/fcell.2021.645593 (2021).

2 He, S. & Sharpless, N. E. Senescence in Health and Disease. Cell 169, 1000–1011, doi:10.1016/j.cell.2017.05.015 (2017).

3 Lyu, G. et al. TGF-beta signaling alters H4K20me3 status via miR-29 and contributes to cellular senescence and cardiac aging. Nat Commun 9, 2560, doi:10.1038/s41467-018-04994-z (2018).

4 Nacarelli, T., Liu, P. & Zhang, R. Epigenetic Basis of Cellular Senescence and Its Implications in Aging. Genes (Basel) 8, doi:10.3390/genes8120343 (2017).

5 Yang, J. H. et al. Loss of epigenetic information as a cause of mammalian aging. Cell 186, 305–326 e327, doi:10.1016/j.cell.2022.12.027 (2023).

6 Crouch, J., Shvedova, M., Thanapaul, R., Botchkarev, V. & Roh, D. Epigenetic Regulation of Cellular Senescence. Cells 11, doi:10.3390/cells11040672 (2022).

7 Kabacik, S. et al. The relationship between epigenetic age and the hallmarks of aging in human cells. Nat Aging 2, 484–493, doi:10.1038/s43587-022-00220-0 (2022).

8 Hubbard, B. P. & Sinclair, D. A. Small molecule SIRT1 activators for the treatment of aging and age-related diseases. Trends in Pharmacological Sciences 35, 146–154, doi:https://doi.org/10.1016/j.tips.2013.12.004 (2014).

9 Vigili de Kreutzenberg, S., et al. Metformin improves putative longevity effectors in peripheral mononuclear cells from subjects with prediabetes. A randomized controlled trial. *Nutrition*, Metabolism and Cardiovascular Diseases 25, 686–693, doi:https://doi.org/10.1016/j.numecd.2015.03.007 (2015).

10 Vaquero, A. & Reinberg, D. Calorie restriction and the exercise of chromatin. Genes Dev 23, 1849–1869, doi:10.1101/gad.1807009 (2009).

11 Li, Y., Daniel, M. & Tollefsbol, T. O. Epigenetic regulation of caloric restriction in aging. BMC Med 9, 98, doi:10.1186/1741-7015-9-98 (2011).

12 Johmura, Y. et al. Senolysis by glutaminolysis inhibition amelioratesvarious age-associated disorders. Science 371, 265–270 (2021).

13 Xu, H., Wu, M., Ma, X., Huang, W. & Xu, Y. Function and Mechanism of Novel Histone Posttranslational Modifications in Health and Disease. Biomed Res Int 2021, 6635225, doi:10.1155/2021/6635225 (2021).

14 Sabari, B. R., Zhang, D., Allis, C. D. & Zhao, Y. Metabolic regulation of gene expression through histone acylations. Nat Rev Mol Cell Biol 18, 90–101, doi:10.1038/nrm.2016.140 (2017).

15 Zhang, D. et al. Metabolic regulation of gene expression by histone lactylation. Nature 574, 575–580, doi:10.1038/s41586-019-1678-1 (2019).

16 Irizarry-Caro, R. A. et al. TLR signaling adapter BCAP regulates inflammatory to reparatory macrophage transition by promoting histone lactylation. Proc Natl Acad Sci U S A 117, 30628–30638, doi:10.1073/pnas.2009778117 (2020).

17 Wang, N. et al. Histone Lactylation Boosts Reparative Gene Activation Post–Myocardial Infarction. Circulation Research 131, 893–908, doi:10.1161/circresaha.122.320488 (2022).

18 Xiong, J. et al. Lactylation-driven METTL3-mediated RNA m6A modification promotes immunosuppression of tumor-infiltrating myeloid cells. Molecular Cell 82, 1660–1677.e1610, doi:10.1016/j.molcel.2022.02.033 (2022).

19 Li, L. et al. Glis1 facilitates induction of pluripotency via an epigenome-metabolome-epigenome signalling cascade. Nat Metab, doi:10.1038/s42255-020-0267-9 (2020).

20 Yu, J. et al. Histone lactylation drives oncogenesis by facilitating m(6)A reader protein YTHDF2 expression in ocular melanoma. Genome Biol 22, 85, doi:10.1186/s13059-021-02308-z (2021).

21 Dai, S.-K. et al. Dynamic profiling and functional interpretation of histone lysine crotonylation and lactylation during neural development. Development 149, doi:10.1242/dev.200049 (2022).

22 Yang, Q. et al. A proteomic atlas of ligand–receptor interactions at the ovine maternal–fetal interface reveals the role of histone lactylation in uterine remodeling. Journal of Biological Chemistry 298, doi:10.1016/j.jbc.2021.101456 (2022).

23 Tian, Q. & Zhou, L.-q. Lactate Activates Germline and Cleavage Embryo Genes in Mouse Embryonic Stem Cells. Cells 11, doi:10.3390/cells11030548 (2022).

24 Pan, R.-Y. et al. Positive feedback regulation of microglial glucose metabolism by histone H4 lysine 12 lactylation in Alzheimer’s disease. Cell Metabolism 34, 634–648.e636, doi:https://doi.org/10.1016/j.cmet.2022.02.013 (2022).

25 Cui, H. et al. Lung Myofibroblasts Promote Macrophage Profibrotic Activity through Lactate-induced Histone Lactylation. Am J Respir Cell Mol Biol 64, 115–125, doi:10.1165/rcmb.2020-0360OC (2021).

26 Moreno-Yruela, C. et al. Class I histone deacetylases (HDAC1-3) are histone lysine delactylases. Sci Adv 8, eabi6696, doi:10.1126/sciadv.abi6696 (2022).

27 Galle, E. et al. H3K18 lactylation marks tissue-specific active enhancers. Genome Biology 23, doi:10.1186/s13059-022-02775-y (2022).

28 Ivashkiv, L. B. The hypoxia–lactate axis tempers inflammation. Nature Reviews Immunology 20, 85–86, doi:10.1038/s41577-019-0259-8 (2019).

29 van Vliet, T. et al. Physiological hypoxia restrains the senescence-associated secretory phenotype via AMPK-mediated mTOR suppression. Mol Cell, doi:10.1016/j.molcel.2021.03.018 (2021).

30 Deng, C. et al. TNFRSF19 Inhibits TGFβ Signaling through Interaction with TGFβ Receptor Type I to Promote Tumorigenesis. Cancer Research 78, 3469–3483, doi:10.1158/0008-5472.CAN-17-3205 (2018).

31 Li, P. et al. Tankyrase Mediates K63-Linked Ubiquitination of JNK to Confer Stress Tolerance and Influence Lifespan in Drosophila. Cell Reports 25, 437–448, doi:10.1016/j.celrep.2018.09.036 (2018).

32 Huang, S.-M. A. et al. Tankyrase inhibition stabilizes axin and antagonizes Wnt signalling. Nature 461, 614–620, doi:10.1038/nature08356 (2009).

33 Silberman, J., Boehlein, J., Abbate, T. & Moore, E. A Biomaterial Model to Assess the Effects of Age in Vascularization. Cells Tissues Organs, doi:10.1159/000523859 (2022).

34 Eklund, L. et al. Lack of type XV collagen causes a skeletal myopathy and cardiovascular defects in mice. Proceedings of the National Academy of Sciences 98, 1194–1199, doi:10.1073/pnas.98.3.1194 (2001).

35 Kim, H. et al. TGF-β2 and collagen play pivotal roles in the spheroid formation and anti-aging of human dermal papilla cells. Aging 13, 19978–19995, doi:10.18632/aging.203419 (2021).

36 Pagliaroli, L. & Trizzino, M. The Evolutionary Conserved SWI/SNF Subunits ARID1A and ARID1B Are Key Modulators of Pluripotency and Cell-Fate Determination. Frontiers in Cell and Developmental Biology 9, doi:10.3389/fcell.2021.643361 (2021).

37 Covarrubias, A. J., Perrone, R., Grozio, A. & Verdin, E. NAD+ metabolism and its roles in cellular processes during ageing. Nature Reviews Molecular Cell Biology 22, 119–141, doi:10.1038/s41580-020-00313-x (2020).

38 Li, J. et al. A conserved NAD+binding pocket that regulates protein-protein interactions during aging. Science 355, 1312–1317, doi:10.1126/science.aad8242 (2017).

39 Dos Santos, M., et al. Single-nucleus RNA-seq and FISH identify coordinated transcriptional activity in mammalian myofibers. Nature Communications 11, doi:10.1038/s41467-020-18789-8 (2020).

40 Angelidis, I. et al. An atlas of the aging lung mapped by single cell transcriptomics and deep tissue proteomics. Nat Commun 10, 963, doi:10.1038/s41467-019-08831-9 (2019).

41 Criswell, T. L. et al. The role of endothelial cells in myofiber differentiation and the vascularization and innervation of bioengineered muscle tissue in vivo. Biomaterials 34, 140–149, doi:10.1016/j.biomaterials.2012.09.045 (2013).

42 Garten, A. et al. Physiological and pathophysiological roles of NAMPT and NAD metabolism. Nat Rev Endocrinol 11, 535–546, doi:10.1038/nrendo.2015.117 (2015).

43 Xie, Y. et al. FGF/FGFR signaling in health and disease. Signal Transduct Target Ther 5, 181, doi:10.1038/s41392-020-00222-7 (2020).

44 Kaur, N. et al. Multi-organ FGF21-FGFR1 signaling in metabolic health and disease. Front Cardiovasc Med 9, 962561, doi:10.3389/fcvm.2022.962561 (2022).

45 Caielli, S. et al. Erythroid mitochondrial retention triggers myeloid-dependent type I interferon in human SLE. Cell 184, 4464–4479 e4419, doi:10.1016/j.cell.2021.07.021 (2021).

46 Hagihara, H. et al. Protein lactylation induced by neural excitation. Cell Rep 37, 109820, doi:10.1016/j.celrep.2021.109820 (2021).

47 Li, M. et al. Oncogene-induced cellular senescence elicits an anti-Warburg effect. Proteomics 13, 2585–2596, doi:10.1002/pmic.201200298 (2013).

48 Unterluggauer, H. et al. Premature senescence of human endothelial cells induced by inhibition of glutaminase. Biogerontology 9, 247–259, doi:10.1007/s10522-008-9134-x (2008).

49 Brooks, G. A. Lactate as a fulcrum of metabolism. Redox Biology 35, doi:10.1016/j.redox.2020.101454 (2020).

50 Shi, Q., Liu, S., Kristiansen, K., Liu, L. & Mathelier, A. The FASTQ+ format and PISA. Bioinformatics 38, 4639–4642, doi:10.1093/bioinformatics/btac562(2022).

51 Petrany, M. J. et al. Single-nucleus RNA-seq identifies transcriptional heterogeneity in multinucleated skeletal myofibers. Nature Communications 11, doi:10.1038/s41467-020-20063-w (2020).

52 Kim, M. et al. Single-nucleus transcriptomics reveals functional compartmentalization in syncytial skeletal muscle cells. Nature Communications 11, doi:10.1038/s41467-020-20064-9 (2020).

53 Jin, S. et al. Inference and analysis of cell-cell communication using CellChat. Nature Communications 12, doi:10.1038/s41467-021-21246-9 (2021).

54 Zhou, Y. et al. Metascape provides a biologist-oriented resource for the analysis of systems-level datasets. Nature Communications 10, doi:10.1038/s41467-019-09234-6 (2019).

55 Gillespie, M. et al. The reactome pathway knowledgebase 2022. Nucleic Acids Research 50, D687–D692, doi:10.1093/nar/gkab1028 (2022).

